# Auto-AVS: A Novel Framework for Automated Autophagic Vacuole Simulation

**DOI:** 10.64898/2026.10.21.760762

**Authors:** Stela Bleta, Payton Dunning, Andrew Ross, Steven K. Backues, Ourania Spantidi

**Affiliations:** Department of Computer Science at Eastern Michigan University, Ypsilanti, USA, 48197; Department of Mathematics and Statistics at Eastern Michigan University, Ypsilanti, USA, 48197; Department of Chemistry at Eastern Michigan University, Ypsilanti, USA, 48197

**Keywords:** Autophagic Vacuole Simulation, Computational Modeling, CompuCell3D, Autophagic Bodies, Autophagy, Stereology

## Abstract

Transmission electron microscopy (TEM) is widely used to quantify autophagic bodies in yeast, but ultra-thin sectioning provides only two-dimensional cross-sections of inherently 3D structures. Inferring the underlying size and number of autophagic bodies therefore requires assumptions about their geometry, spatial organization, and distribution within the vacuole. We introduce Auto-AVS, an open-source computational stereology framework that combines configurable 3D autophagic-vacuole simulation, optional CompuCell3D-based body deformation, virtual TEM sectioning, and statistical comparison with experimental data. We first validate the virtual sectioning pipeline against analytical stereology and an existing spherical R-based simulation. We then use Auto-AVS to examine whether explicitly modeling autophagic-body deformation alters observed section statistics and find no consistent measurable effect under the tested simulation parameters. Finally, we apply the framework to experimental wild-type and mutant TEM datasets. Auto-AVS produces body-size estimates closely consistent with conventional stereological estimates, while body-number fitting reveals substantial discrepancies between simulated and experimental distributions despite similar average count estimates. These discrepancies implicate spatial clustering and vacuole geometry as important limitations of current body-number models. Rather than replacing conventional stereology, Auto-AVS provides a computational framework for testing its underlying assumptions and determining which aspects of 3D organization affect inference from two-dimensional microscopy.

## I. Introduction

Macroautophagy (hereafter simply referred to as autophagy) is a subcellular stress response pathway shared among animals, plants, and fungi. It involves wrapping cytoplasmic cargo in a double-membraned vesicle termed an autophagosome for delivery to the lysosome (in animals) or vacuole (in plants and fungi). Here the cargo is degraded and the nutrients recycled. Autophagy is important both for maintaining cellular energy balance and for preserving cellular health by getting rid of damaged or malfunctioning cellular components [1]. In humans, it plays a particular role in preventing neurodegenerative diseases such as Parkinson’s and Alzheimer’s [2]. For this reason, there is extensive research ongoing into how this process works and in particular the roles played by various autophagy proteins.

The central event in autophagy is formation of the autophagosome. One way to understand the roles of the proteins involved in autophagosome formation is to ask whether each of them controls autophagosome size or autophagosome number. This research can be done in baker’s yeast, (*S. cerevisiae*), a commonly used model organism for studying autophagy. Here, it has been shown that the protein Atg8 controls the size of autophagosomes [3], whereas the protein Atg9 controls autophagosome number [4]. Atg7 controls both size and number [5], and research into additional proteins is ongoing.

The autophagosome is a transient structure: almost as soon as it is formed, it fuses with the vacuole membrane, delivering the inner membrane and the cargo into the vacuole for degradation. Therefore, we measure the size and number of autophagosomes formed via a proxy: autophagic bodies. These are the remnants of the autophagosomes after fusion with the vacuolar membrane: the inner autophagosomal membrane plus the cargo that has been delivered into the vacuole. Under normal conditions these are rapidly degraded, but in yeast mutants that lack protease activity, they are stabilized. When these mutants are starved for a few hours to induce autophagosome formation, dozens of autophagic bodies accumulate in the vacuole where they can be counted and measured via transmission electron microscopy (TEM) [6].

### Motivation

Although TEM is the gold standard for visualizing autophagic bodies, it only captures an image of a thin slice of a cell - only 70 nm thick, compared to a total cell size of around 10,000 nm. Therefore, it does not give a view of the entire autophagic bodies, but instead only a random sampling of body cross-sections. This sectioning inherently biases both size and count measurements: larger bodies produce larger cross-sectional areas (and are intersected more often), while smaller or sparse bodies can be entirely missed. As a result, raw cross-section distributions cannot be directly equated to the true 3D dimensions or abundance of autophagic bodies. However, there will be a statistical relationship between the original bodies and their cross-sections: if the bodies are larger, then the cross-sections will also be larger, on average, and if there are more bodies, then on average more cross-sections will be shown [6], [7].

The field of *stereology* formalizes exactly this challenge of inferring three-dimensional structure from two-dimensional slices. In its classical form (often referred to as Wicksell’s corpuscle problem [8]), one seeks the true size distribution of spherical particles given only their planar cross-section measurements. However, Wicksell’s solution assumes that all bodies are perfect spheres and are randomly dispersed without overlap. These are assumptions that do not strictly hold for autophagic bodies, which can vary in shape and exhibit spatial clustering.

To investigate these limitations, we introduce Auto-AVS, a computational stereology framework that explicitly models the 3D organization of autophagic bodies and reproduces the thin-section sampling process used in TEM. Unlike closed-form stereological approaches, Auto-AVS allows assumptions about body geometry, deformation, spatial packing, and vacuole organization to be modified and evaluated directly. The framework can therefore be used not only to estimate underlying 3D parameters from experimental cross-sections, but also to determine which modeling assumptions meaningfully influence those estimates. Importantly, agreement in a final point estimate does not necessarily imply that the underlying simulated cross-section distribution faithfully reproduces the experimental observations. Auto-AVS therefore evaluates both inferred parameters and the full distributions from which those estimates arise.

The contributions of this work are:

- We develop a configurable Python-based 3D simulation and virtual TEM-sectioning framework for modeling autophagic bodies within yeast vacuoles and measuring the resulting two-dimensional cross-sections.
- We integrate an optional CompuCell3D-based [9] deformation stage that enables direct testing of whether deviations from spherical body geometry measurably affect stereological observations.
- We systematically compare simulation-derived section statistics with analytical stereology and a previously developed spherical R-based simulation, characterizing where Auto-AVS reproduces conventional spherical predictions and where increasing body density and size produce divergent count behavior.
- We apply Auto-AVS to experimental wild-type and mutant TEM datasets and demonstrate that body-size estimates are consistent with conventional stereological estimates, while full-distribution analysis exposes limitations in body-number reconstruction that are not apparent from average estimates alone.
- Through these experiments, we show that body-number inference is substantially more sensitive than body-size inference to assumptions about spatial organization and vacuole geometry, and demonstrate how simulation-based stereology can expose limitations that are hidden by agreement in summary estimates alone.

Auto-AVS is released as open-source software alongside instructions for reuse and reproducibility.^1^

## II. Related Works

Early efforts for estimating autophagic-body size and count relied on closed-form stereology and relatively simple computational tools. The work in [6] implemented an Excel-based approach that applies Wicksell’s corpuscle formula to cross-sectional counts and sizes, assuming spherical bodies to infer their underlying 3D size and number. Related work in [5] and [7] used R-based simulations with sampled body radii and counts, but likewise modeled autophagic bodies as spheres. Such an assumption can become problematic when the underlying structures are irregular or deformed, since object shape affects the probability and geometry of intersection with a planar section.

The mathematical basis of these approaches derives from Wicksell’s corpuscle problem [8], which provides a framework for recovering the size distribution of randomly dispersed spherical objects from planar sections. However, biological organelles frequently exhibit complex three-dimensional morphologies that are not well represented by idealized spheres. Recent advances in volumetric electron microscopy, including focused ion beam scanning electron microscopy (FIB-SEM) and serial block-face SEM, have therefore increasingly been used to reconstruct and quantify organelles directly in three dimensions [10], [11]. For example, Müller et al. [10] reconstructed multiple intracellular organelle populations across entire pancreatic *β* cells using FIB-SEM, while Garza-Lopez et al. [11] presented a workflow for extracting quantitative 3D organelle morphology from volumetric electron-microscopy data. More recently, Barad et al. [12] developed a surface-morphometrics pipeline for cryo-electron tomography that explicitly quantifies complex and pleomorphic organelle membrane geometries. These studies demonstrate the value of three-dimensional representations when morphology departs from simple geometric assumptions. However, obtaining complete volumetric EM reconstructions requires specialized imaging and substantial segmentation and analysis effort.

Computational spatial models offer a complementary approach in which three-dimensional biological structures can be represented explicitly and then interrogated in silico. Cellular Potts and related spatial modeling frameworks have been used to reproduce experimentally observed cellular geometries and mechanical behaviors, including irregular and deformable shapes [9], [13]. In particular, Nürnberg et al. [13] demonstrated a recent pipeline for constructing Cellular Potts simulations from experimentally derived three-dimensional biological geometries, illustrating how image-derived morphology and mechanistic spatial simulation can be combined.

Our work builds on these two lines of research. Rather than analytically correcting 2D measurements under an assumed spherical geometry or requiring direct volumetric reconstruction of every experimental specimen, Auto-AVS generates explicit 3D populations of autophagic bodies, applies virtual TEM sectioning, and compares the resulting 2D observations with experimental TEM distributions. This enables assumptions concerning size, number, spatial arrangement, sectioning, vacuole geometry, and non-spherical morphology to be evaluated within the same computational framework rather than being fixed a priori.

## III. Methodology

### A. Simulation Pipeline

The Auto-AVS pipeline consists of four main stages:

i. specification of user-defined simulation parameters,
ii. generation and placement of spherical autophagic bodies,
iii. optional CompuCell3D-based deformation, and (iv) virtual TEM sectioning and cross-section measurement. The resulting measurements are aggregated across repeated simulation runs and exported for downstream statistical analysis. The modular design allows individual modeling assumptions, including body size, body count, spatial packing, deformation, and sectioning parameters, to be varied independently.

#### 1) User-defined Input Parameters

Simulation parameters are specified through a configuration file, enabling reproducible experiments and systematic exploration of alternative parameter settings. Parameters not explicitly provided by the user are assigned default values. The principal configurable parameters are:

- **Sample size:** The number of virtual vacuoles generated and sectioned for each parameter configuration. This value is selected according to the experimental design.
- **Scale factor:** A conversion factor, expressed in nm per voxel, that determines the spatial resolution of the simulation. Increasing the scale factor decreases computational cost at the expense of spatial resolution.
- **Vacuole radius distribution parameters (***µ* **and** *σ*): Distribution parameters of the log-normal distribution used to generate the inner vacuole radius. The outer radius is defined as 5% larger than the inner radius to represent the vacuolar boundary. A log-normal distribution was used to ensure positive-valued radii while allowing right-skewed size variation.
- **Body radius distribution parameters (***µ* **and** *σ*): Parameters of the log-normal distribution used to generate autophagic-body radii. Auto-AVS can iterate over a user-defined range of mean values to evaluate alternative underlying size distributions.
- **Body number distribution parameters (***µ* **and** *σ*): Parameters governing the distribution of the number of autophagic bodies generated within each simulated vacuole. The framework supports evaluation over a user-defined range of mean body counts.
- **TEM parameters:** Parameters controlling virtual section generation and detection, including section thickness, minimum visible vacuole radius, and minimum detectable body-section size.

#### 2) Generating Spheres

Auto-AVS initially represents autophagic bodies as spheres before any optional deformation is applied. The sphere-generation stage constructs a configurable three-dimensional population of bodies within a vacuole while accounting for variation in body size, body number, and spatial packing. Generation proceeds in three stages.

##### Step 1 – Number and radius generation

For each simulated cell, the number of autophagic bodies (APBs) is sampled from the specified log-normal body-count distribution. The radius of each APB is independently sampled from the specified log-normal body-radius distribution. The sampled radii are retained in their generated order rather than being sorted by size prior to placement.

##### Step 2 – Placement of bodies

The generated APBs are placed so as to form a compact, non-overlapping cluster. Placement is initialized using direction vectors derived from a hexagonal close-packed (HCP) arrangement of equal-sized spheres with randomized initial orientation. The first APB is placed at the origin. Each subsequent APB is assigned a direction vector, and candidate positions are evaluated incrementally along that direction until a location is found that does not intersect any previously placed body.

Because this procedure is greedy and does not modify earlier placements, the resulting configuration is not guaranteed to be maximally compact, though, actual yeast cells also might not maximize compactness. Auto-AVS therefore provides an optional nonlinear optimization stage that minimizes the sum of APB distances from the origin subject to non-overlap constraints. The maximum number of optimization iterations is user-defined; setting this value to zero disables the optimization stage. This procedure is conceptually related to the packing optimization used in [6].

##### Step 3 – Vacuole radius generation

Once the APB positions have been established, a candidate vacuole radius is sampled from the specified log-normal distribution. The vacuole is centered at the origin, coincident with the center of the APB cluster. If the sampled radius is insufficient to enclose all generated APBs without intersection with the vacuolar boundary, a new radius is sampled. The position of a slice through the vacuole is then randomly determined, and checked to ensure that the radius of the spherical cap generated by the slice is above the user-defined vacuole minimum recognition limit. This accounts for the fact that larger vacuoles are more likely to be captured in a TEM section. This process is repeated until a feasible vacuole is obtained. After 100 failed tries, a vacuole just large enough to hold the bodies is generated instead. The number of rejected candidate radii is recorded because repeated rejection of smaller radii can shift the realized vacuole-size distribution away from the specified input distribution.

After a feasible vacuole is generated, the position of the virtual TEM section through the vacuole is randomly selected. The resulting vacuolar cross-section is checked to ensure that its radius exceeds the user-defined minimum vacuole-recognition threshold. Sections below this threshold are rejected and resampled. This procedure accounts for the greater probability that larger vacuoles will be represented in experimentally observable TEM sections.

The HCP-based procedure provides a computationally efficient means of generating compact, non-overlapping APB configurations, while the optional optimization stage permits tighter packing when additional computational cost is acceptable. In the current implementation, the APB cluster and vacuole are both centered at the origin. This provides a controlled baseline spatial configuration Alternative arrangements, including off-center or membrane-proximal clustering, can be incorporated in future extensions of the framework.

#### 3) Integrating CompuCell3D

Following sphere generation, each simulated vacuole can optionally be passed through a CompuCell3D deformation stage. Auto-AVS first converts the generated geometry into a voxel-based representation compatible with CompuCell3D. Each occupied voxel is stored individually together with information identifying whether it belongs to an APB or to the vacuolar boundary and its corresponding spatial coordinates. This representation preserves the discretized three-dimensional geometry generated during the sphere-placement stage and is used to initialize the subsequent CompuCell3D simulation.

The resulting representation is used to initialize CompuCell3D automatically. Cell-type assignments, mechanical constraints, Monte Carlo schedules, and plugin configurations are generated programmatically, allowing simulations to be executed in headless batch mode without manual GUI interaction or configuration-file editing. During this stage, CompuCell3D applies the specified mechanical interactions to deform the initially spherical APBs into non-spherical configurations. Figure 2 illustrates an example configuration over 500 simulation steps.

**Fig. 1.**
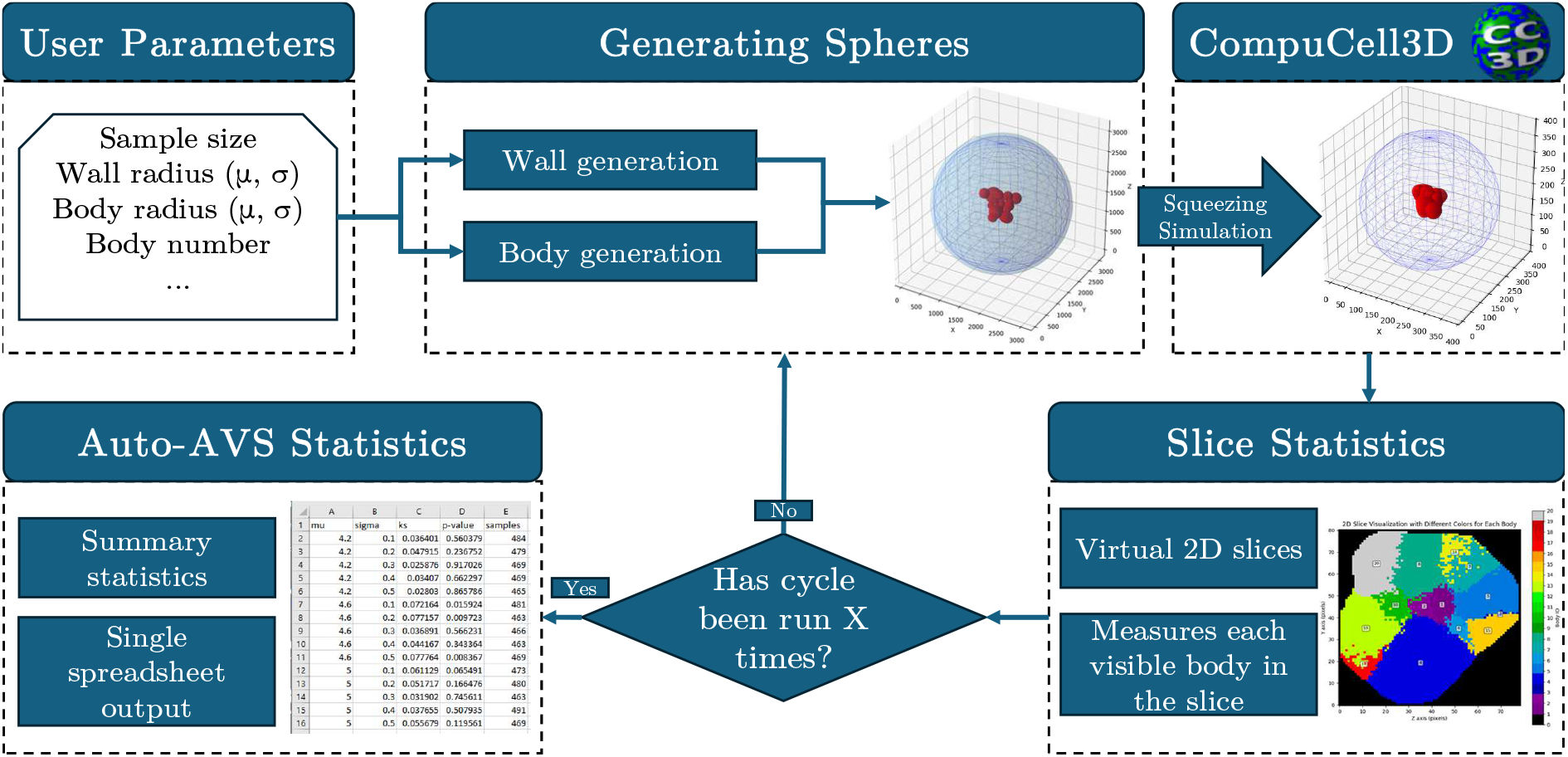
An overview of the Auto-AVS simulation pipeline. User-defined parameters control generation of spherical autophagic bodies and vacuole geometry, CompuCell3D deformation, virtual 2D sectioning, slice statistics, repeated simulation cycles, and final aggregated Auto-AVS statistics.

**Fig. 2.**
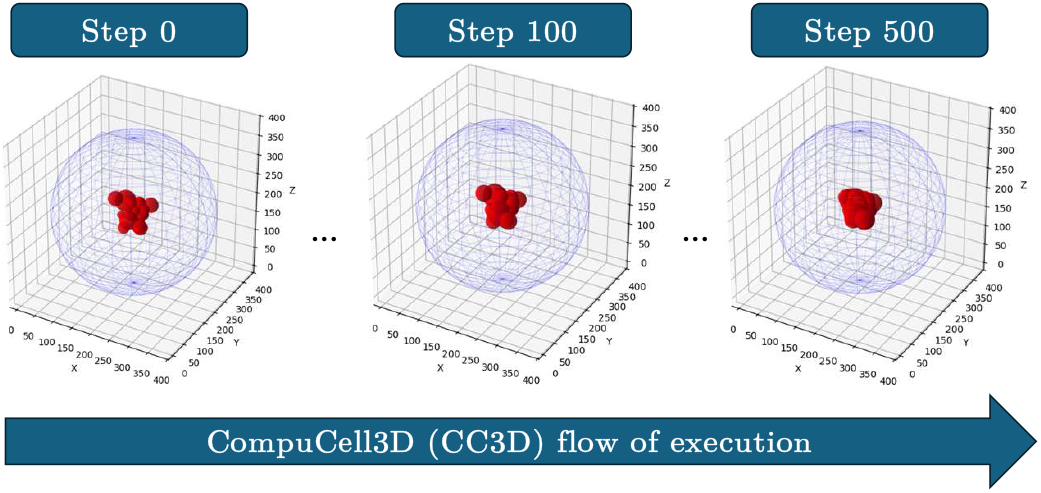
Example CompuCell3D deformation sequence for simulated autophagic bodies within a vacuole, as described in Section III-A2. Three 3D snapshots at simulation steps 0, 100, and 500 show the red body cluster changing shape over time inside the translucent vacuolar boundary.

Because the CompuCell3D stage is optional, otherwise identical simulated populations can be evaluated both before and after deformation. This enables the effect of non-spherical body geometry on subsequent virtual TEM measurements to be isolated directly. The present configuration is intended as a controlled deformation model rather than a claim that the resulting APB morphology fully reproduces the morphology observed experimentally.

#### 4) Slice-based Cross-section Measurements

To emulate TEM sectioning, Auto-AVS samples each three-dimensional simulation with an *x*-aligned slab of user-defined physical thickness. The specified thickness is converted to lattice units using the simulation-to-physical scale factor. The slab position is selected according to a configurable policy, with typical usage being to use the slice position defined during the vacuole generation step, as this most accurately reproduces the random sampling from actual TEM images.

For each APB intersecting the slab, all APB-labeled voxels within the section are retained and projected onto the *y*–*z* plane to produce a two-dimensional binary cross-section mask. Cross-sections are then quantified using connected-component morphometrics, including area, perimeter, and major- and minor-axis lengths. Components smaller than a user-defined detectability threshold, derived from the minimum body-section size, are excluded from subsequent analysis.

If a single three-dimensional APB produces multiple spatially disconnected above-threshold components within the same section, each component is treated as an independent detected cross-section for slice-level statistics. All measurements are converted from lattice units to physical units and recorded for aggregation across replicate simulations and parameter settings.

For comparison with analytical stereology, each measured cross-sectional area *A* is additionally expressed as an equivalent circular radius,

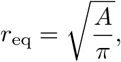

which corresponds to the radius of a circle having the same measured area as the detected cross-section.

These stages are orchestrated automatically so that large parameter sweeps, repeated simulation runs, CompuCell3D execution, virtual sectioning, and downstream statistical aggregation can be performed without manual reconfiguration between runs.

#### 5) Statistical Comparison of Real and Simulated Data

Once measurements have been recorded from slices of many simulated vacuoles, the distributions of APB cross-sectional radii and number of APB cross-sections per slice can be compared to data derived from actual TEM data [6] to generate estimates of the original size and number distribution of ABPs in the biological sample. We provide a set of tools for this comparison that include calculation of summary statistics, goodness-of-fit comparisons of the distributions of derived from real data to that derived from simulated data runs with different parameters, estimations of optimal values of mu and sigma based on linear or polynomial regression, and various options for graphical comparison of the real and simulated data.

### B. Statistical Analysis and Validation

To validate the Auto-AVS virtual sectioning and measurement pipeline, we compare its outputs against both analytical stereological predictions and an existing R-based simulator that assumes spherical autophagic bodies [5]. The purpose of this validation is to determine whether Auto-AVS reproduces the expected behavior of conventional stereology under conditions in which the underlying geometric assumptions are satisfied.

Simulations are conducted across a range of log-normal body-size distributions defined by parameters *µ* and *σ*, and for multiple body-count configurations. In the current validation experiments, we consider populations containing 1, 20, and 40 bodies per vacuole. For each configuration, we measure two quantities derived from the virtual TEM sections:

1. the fraction of generated autophagic bodies intersected by at least one virtual section, and
2. the mean equivalent cross-section radius observed in the resulting two-dimensional sections.

The same quantities are computed using the spherical R-based simulator and analytical stereological calculations, enabling direct comparison across methods. Under spherical assumptions, agreement among these three approaches provides evidence that the Auto-AVS geometry generation, virtual sectioning, and cross-section measurement procedures behave as expected.

The resulting validation experiments establish a baseline for the subsequent analyses. In particular, they allow deviations introduced by additional modeling assumptions, including non-spherical deformation and spatial organization, to be evaluated relative to the known spherical case.

### C. Evaluation of CompuCell3D-Based Body Deformation

To determine whether explicitly modeling non-spherical autophagic-body geometry alters the measurements obtained from virtual TEM sections, we compared Auto-AVS simulations performed with and without the CompuCell3D deformation stage. In the spherical condition, generated APBs were passed directly to the virtual sectioning stage. In the deformation condition, APBs generated from the same input distributions were first processed for 500 CompuCell3D simulation steps using the contact-based deformation configuration and were then sectioned using the same virtual TEM procedure.

The comparison was performed across multiple combinations of APB size and number distributions in order to determine whether the effect of deformation depended on the characteristics of the simulated population. The tested configurations included small, medium, and larger body-size distributions together with small, intermediate, and large body-count distributions. For each configuration, replicate vacuoles were generated and analyzed under both spherical and CompuCell3D-deformed conditions. Most deformation experiments consisted of 200 simulated vacuoles per configuration. For the small-body, small-count configuration, the experiment was repeated independently, and four CompuCell3D runs were additionally pooled to obtain a larger sample of 800 simulated vacuoles.

Two primary outcomes were used to evaluate the effect of deformation: (i) the size of the APB cross-sections detected in virtual TEM sections and (ii) the number of APB cross-sections detected per vacuole section. The fraction of generated APBs intersected by the virtual section was also recorded as a descriptive measure of sectioning probability. Mean cross-section size and mean number of detected cross-sections were compared between the spherical and deformed conditions using two-sample *t*-tests.

This analysis was designed as a sensitivity test of the spherical-body assumption. Rather than assuming that explicit deformation necessarily improves stereological inference, the objective was to determine whether the degree of deformation produced by the current CompuCell3D configuration is sufficient to measurably alter the two-dimensional quantities observable in TEM sections.

### D. Simulation-Based Fitting to Experimental TEM Data

Auto-AVS was subsequently used to estimate the underlying three-dimensional size and number distributions of autophagic bodies from experimental TEM data. Two experimental datasets were analyzed independently: a wild-type (WT) sample and a mutant sample exhibiting smaller and fewer observed body cross-sections. The fitting procedure was performed sequentially, with body size estimated before body number because the probability of observing a body in a TEM section depends on its size, whereas the estimated body-size distribution is insensitive to the total number of bodies present.

#### 1) Body-Size Fitting

Body-size fitting was performed by comparing experimentally observed cross-section sizes with simulated cross-section distributions generated over a grid of candidate log-normal body-radius parameters (*µ, σ*). The initial simulation library covered *µ* values from 4.6 to 5.2 and *σ* values from 0.2 to 0.4. Each parameter combination was simulated using 40 APBs per vacuole and 10,000 virtual vacuoles. These fitting simulations were performed without CompuCell3D deformation in order to isolate the effects of the underlying size distribution.

Candidate parameter combinations were evaluated using several complementary statistics implemented in AVSStats. First, the mean and standard deviation of the simulated cross-section-size distribution were calculated for every (*µ, σ*) combination. Regression surfaces relating the input *µ* and *σ* values to these two observed statistics were then used to estimate the parameter combination expected to reproduce the mean and standard deviation of the experimental data.

Because matching only the first two moments does not ensure agreement between complete distributions, we additionally used *AVSstats* to compare each simulated cross-section distribution with the corresponding experimental distribution using the two-sample Kolmogorov–Smirnov (KS) and Epps– Singleton (ES) statistics. For both measures, smaller values indicate greater similarity between the simulated and experimental distributions. The resulting statistics were evaluated directly over the simulation grid and were also fit as second-order polynomial functions of *µ* and *σ* to estimate parameter values associated with the minimum KS and ES statistics.

The fitting procedure followed a coarse-to-fine strategy. After the initial parameter sweep, additional simulations were generated over narrower ranges surrounding the best-supported region. For the WT dataset, the refined search considered *µ* values from 5.0 to 5.15 and *σ* values from 0.15 to 0.30. For the mutant dataset, the refined search considered *µ* values from 4.65 to 4.80 and *σ* values from 0.15 to 0.30.

Final candidate fits were assessed using both the quantitative comparison statistics and graphical diagnostics generated by AVSStats. These included cumulative distribution function (CDF) plots and ridgeline plots for comparing multiple simulation configurations, as well as quantile–quantile (Q–Q) plots and violin plots for detailed comparison of selected simulated and experimental distributions. Parameter estimates were selected by considering agreement across these complementary measures rather than relying on a single goodness-of-fit statistic.

#### 2) Body-Number Fitting

After estimating the body-size distribution for each experimental dataset, these size parameters were held fixed while the underlying body-number distribution was varied. Candidate APB counts were modeled using log-normal distributions parameterized by body-number *µ* and *σ*. The simulated number of APB cross-sections observed in each virtual vacuole section was then compared with the experimentally observed number of cross-sections per vacuole section.

The fitting procedure paralleled that used for body size. For each candidate body-number distribution, AVSStats calculated the mean and standard deviation of the number of detected APB cross-sections. Regression models relating the body-number parameters (*µ, σ*) to these summary statistics were used to estimate parameter combinations consistent with the experimental observations. Complete count distributions were additionally compared using the KS and ES statistics, and second-order polynomial fits to these statistics were used to identify candidate regions of improved distributional agreement. CDF, Q–Q, violin, and ridgeline plots were used to examine differences in distribution shape that were not captured by the summary statistics alone.

For the WT dataset, simulations were performed using the fitted WT body-size parameters while varying body-number *µ* from 3.7 to 4.8 and *σ* from 0 to 0.4. For the mutant dataset, the simulations used the mutant body-size parameters, and additionally used the experimentally determined mutant vacuole-size distribution (*µ* = 6.172, *σ* = 0.417), rather than the WT vacuole-size distribution (*µ* = 7.101, *σ* = 0.265), to account for the smaller vacuoles observed experimentally.

Because the mutant exhibited substantially fewer observed APB cross-sections, its body-number search was performed iteratively over a lower parameter range. An initial sweep varied body-number *µ* while holding *σ* = 0. A subsequent set of simulations evaluated *µ* values approximately between 1.9 and 2.6 while varying *σ* from 0 to 0.4. When the resulting fits suggested that greater body-number variability was required, the search was expanded to include *σ* values as large as 1.2.

For both WT and mutant datasets, the final body-number estimate was selected by jointly considering agreement in the mean and standard deviation, the KS and ES distribution-comparison statistics, and the corresponding graphical diagnostics. This distribution-level evaluation also allowed discrepancies between simulated and experimental count distributions to be identified even when their average numbers of observed cross-sections were similar.

## IV. Results

### A. Validation against analytical stereology and R simulation

We first compared Auto-AVS against analytical stereological calculations and the previously developed spherical R-based simulation [5] across body populations of 1, 20, and 40 APBs per vacuole. Figure 3 summarizes the percentage difference of each simulation method relative to the analytical calculation for both equivalent cross-section radius and the fraction of generated bodies intersected by the virtual section.

**Fig. 3.**
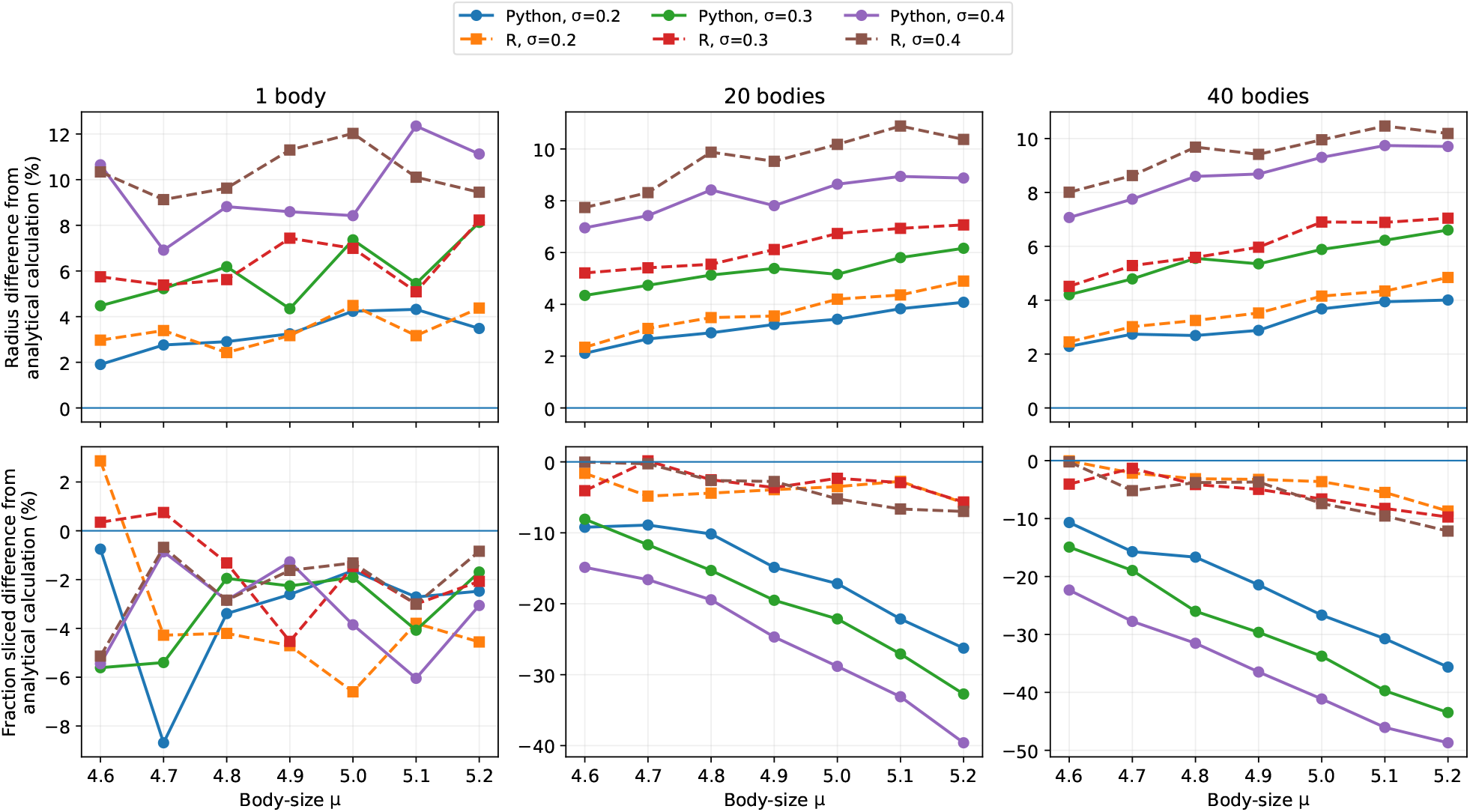
Comparison of Auto-AVS and the previously developed R-based spherical simulation against analytical stereological predictions [5]. Percentage differences are shown for equivalent cross-section radius (top row) and the fraction of generated bodies intersected by the virtual section (bottom row) across populations of 1, 20, and 40 bodies per vacuole. Cross-sectional radius estimates remain comparatively stable across methods, whereas Auto-AVS increasingly diverges from the analytical prediction for the fraction of bodies observed as body count, body-size mean *µ*, and body-size variability *σ* increase.

For equivalent cross-section radius, Auto-AVS and the R-based simulation exhibited similar behavior across all three body-count conditions. Differences relative to the analytical calculation remained comparatively modest and did not increase substantially as the number of bodies per vacuole increased. Instead, the discrepancy was primarily associated with the variability of the underlying body-size distribution: larger values of *σ* generally produced greater differences from the analytical estimate. Across the tested conditions, Auto-AVS and the R-based simulation therefore remained broadly consistent in their estimates of observed cross-sectional body size.

A different pattern was observed for the fraction of generated bodies intersected by the virtual section. With a single body per vacuole, both simulation approaches remained relatively close to the analytical prediction, with no strong systematic dependence on body-size *µ*. As the number of bodies increased, however, Auto-AVS increasingly predicted fewer intersected bodies than the analytical calculation. This divergence became progressively larger with increasing body-size *µ* and *σ* and was most pronounced in the 40-body condition, where the difference approached 50% for the largest and most variable body-size distributions tested. In contrast, the R-based simulation remained substantially closer to the analytical prediction across the same configurations.

These results indicate that the virtual sectioning and measurement procedures produce comparatively stable cross-sectional size estimates across methods, while the probability of observing bodies becomes increasingly sensitive to the representation of higher-density body populations. The divergence therefore emerges primarily in body-count-related behavior rather than in the measurement of the sizes of detected cross-sections.

### B. Effect of CompuCell3D deformation

We next evaluated whether explicitly deforming the initially spherical APBs using CompuCell3D measurably altered the size or number of cross-sections observed in virtual TEM sections. Table I reports the percentage change between the spherical and CompuCell3D-deformed conditions together with the corresponding two-sample *t*-test results.

**TABLE I.** Effect of CompuCell3D deformation on virtual tem measurements. Values report the percentage change relative to the corresponding spherical simulation.

| Configuration | $\Delta$ Size (%) | $p$ | $\Delta$ Count (%) | $p$ |
| --- | --- | --- | --- | --- |
| Medium size, medium count | -2.1 | 0.096 | +1.0 | 0.899 |
| Medium size, large count | -0.7 | 0.563 | -5.6 | 0.462 |
| Medium size, small count | -0.5 | 0.759 | +5.3 | 0.538 |
| Small size/count, run 1 | +4.8 | 0.025 | +15.3 | 0.159 |
| Small size/count, run 2 | +2.9 | 0.160 | +16.6 | 0.128 |
| Small size/count, run 3 | -0.2 | 0.917 | +5.0 | 0.648 |
| Small size/count, run 4 | -1.8 | 0.437 | -6.8 | 0.530 |
| Small size/count, pooled | +1.6 | 0.152 | +7.5 | 0.184 |
| Small size, large count | -0.4 | 0.727 | +0.4 | 0.963 |
| Large/variable size, $\sim 40$ bodies | -2.2 | 0.528 | -7.2 | 0.365 |

Note: For size comparisons, $n$ denotes the number of APB cross-sections; for count comparisons, $n$ denotes the number of simulated vacuoles. Individual CC3D runs used 200 vacuoles and the pooled condition used 800; corresponding spherical simulations used approximately 9,400–10,000 vacuoles.

Across the tested body-size and body-count configurations, CompuCell3D deformation did not produce a consistent change in mean observed cross-section size. Most individual comparisons showed small positive or negative differences and were not statistically significant. One small-body, small-count experiment initially produced a nominally significant increase in observed cross-section size of 4.8% (*p* = 0.025). However, this effect was not reproduced in three independent repetitions, which produced changes of approximately +2.9%, −0.2%, and −1.8%, respectively. When the four runs were pooled, the observed size difference decreased to 1.6% and was not statistically significant (*p* = 0.152).

The number of detected APB cross-sections showed a similarly inconsistent response to deformation. Individual configurations produced both increases and decreases relative to the spherical condition, including comparatively large positive differences in two of the small-body, small-count runs. None of the count comparisons, however, reached statistical significance. The pooled small-body, small-count experiment showed a 7.5% increase in detected cross-sections (*p* = 0.184), while the large and highly variable body-size condition showed a 7.2% decrease (*p* = 0.365).

Taken together, these experiments provide no evidence that the degree of deformation produced by the current CompuCell3D configuration consistently changes either of the section-level quantities used for inference. Thus, although CompuCell3D produces visibly non-spherical APB geometries, the resulting deformation was not sufficient under the tested conditions to produce a reproducible measurable effect on observed cross-section size or number.

### C. Fitting Autophagic-Body Size to Experimental TEM Data

We next evaluated whether Auto-AVS could recover plausible underlying body-size distributions from experimental TEM measurements. The fitting procedure was applied independently to the WT and mutant datasets using the coarse-to-fine parameter search described in Section III-D.

For the WT dataset, the initial search localized the best-fitting region to body-size *µ* values near 5.0–5.1 and *σ* values near 0.2. Refinement over a narrower parameter range produced closely agreeing estimates across the different fitting criteria. Regression against the observed mean and standard deviation estimated *µ* = 5.08 and *σ* = 0.191, while the minimum KS and ES statistics occurred at or near *µ* = 5.05, *σ* = 0.20. Considering the distribution-level comparisons together with these summary statistics, we selected *µ* = 5.05, *σ* = 0.20 as the final WT body-size estimate.

The mutant dataset exhibited substantially smaller observed cross-sections. After refinement, the regression, KS, and ES criteria similarly converged on a comparatively narrow region of parameter space. The final selected body-size distribution was *µ* = 4.75, *σ* = 0.25. Figure 4 compares the experimental size distributions with virtual TEM measurements produced using the selected WT and mutant parameter sets. In both datasets, the simulated distributions closely reproduce the central portion and overall shape of the experimental distributions, with comparatively small deviations confined primarily to the extreme upper tail.

**Fig. 4.**
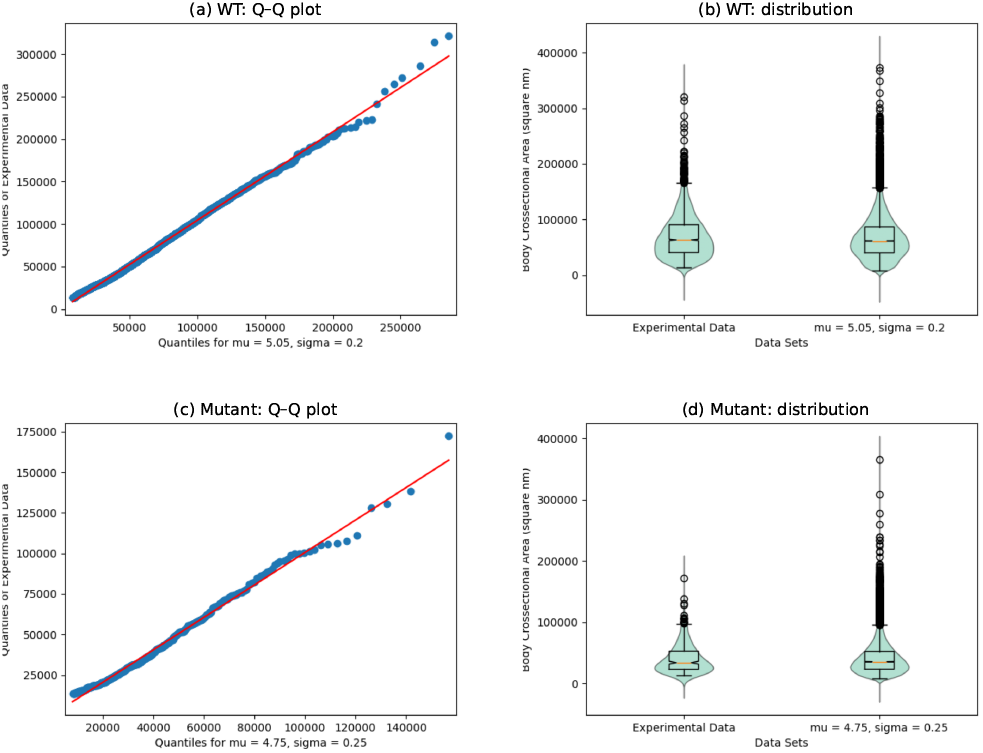
Comparison of experimentally observed and simulated autophagic-body cross-section sizes for the final Auto-AVS body-size fits. (a–b) WT experimental data compared with the selected simulation using *µ* = 5.05 and *σ* = 0.20. (c–d) Mutant experimental data compared with the selected simulation using *µ* = 4.75 and *σ* = 0.25. Q–Q and violin plots show close agreement across most of the observed distributions, supporting comparatively robust inference of underlying body size.

The resulting inferred mean body sizes were 159 nm for WT and 120 nm for the mutant. These values closely agree with the corresponding conventional stereological estimates of 164 nm and 120 nm, respectively (Table II). Thus, despite differences in how the two methods reconstruct the underlying three-dimensional population, body-size inference was comparatively stable across the two approaches.

**TABLE II.**
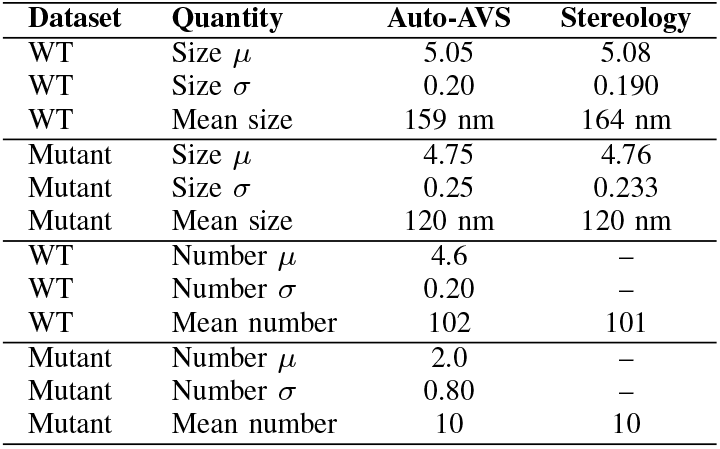
Final Auto-AVS parameter estimates and comparison with conventional stereological estimates. A dash indicates that the corresponding log-normal parameter is not estimated by the conventional stereological method.

### D. Fitting Autophagic-Body Number to Experimental TEM Data

Body-number fitting produced a substantially different result. For the WT dataset, the experimental sections contained an average of 13.0 detected body cross-sections with a standard deviation of 10.6. Although these summary statistics fell within the range produced by the candidate simulations, no tested body-number distribution reproduced the complete experimental count distribution closely.

The best overall compromise was obtained using body-number parameters *µ* = 4.6 and *σ* = 0.20. However, Figure 5 shows that substantial distributional differences remained. In particular, the simulated data produced more vacuole sections containing no visible bodies than were observed experimentally. At the same time, the simulated distribution contained a heavier upper tail, with some sections containing substantially more detected bodies than expected from the experimental data. Consequently, changes in *µ* and *σ* shifted the location and spread of the simulated count distribution but did not eliminate the characteristic mismatch in its shape.

**Fig. 5.**
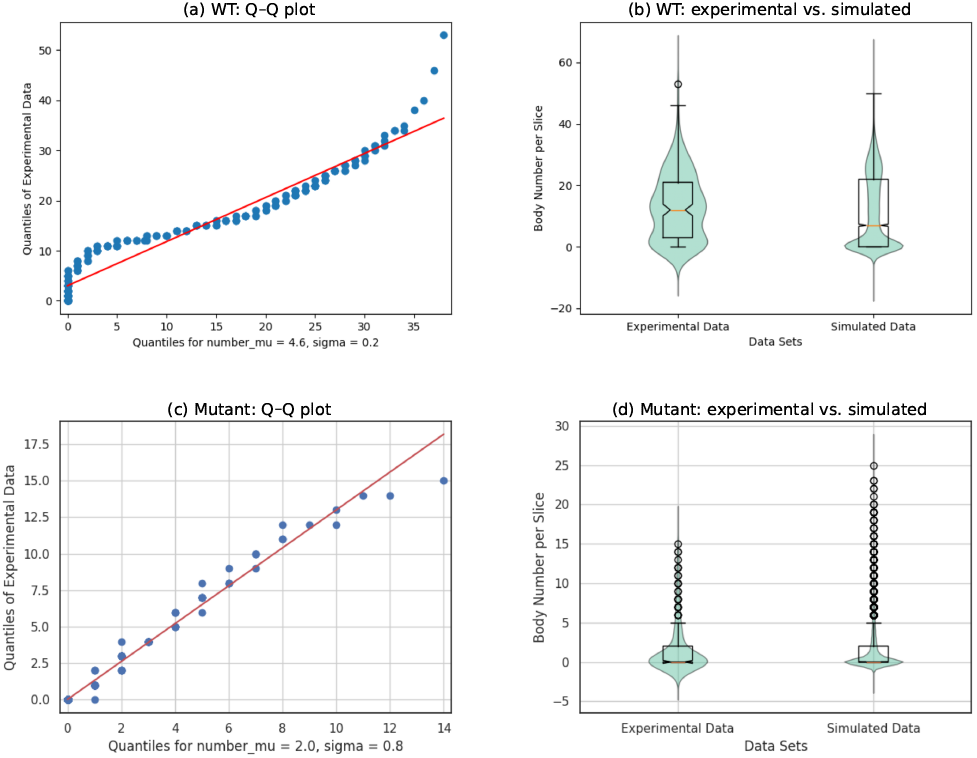
Comparison of experimentally observed and simulated numbers of autophagic-body cross-sections for the selected Auto-AVS body-number fits. (a–b) WT experimental data compared with the selected simulation using body-number *µ* = 4.6 and *σ* = 0.20. (c–d) Mutant experimental data compared with the selected simulation using *µ* = 2.0 and *σ* = 0.80. In contrast to the body-size fits, the simulated count distributions retain substantial shape differences from the experimental data, including excess low-count observations and heavy upper tails, particularly in the WT fit.

The mutant dataset contained substantially fewer observed body cross-sections, with a mean of 1.7 per vacuole section, and also exhibited a smaller experimental vacuole-size distribution. Fitting therefore required exploration of lower body-number *µ* values and substantially larger values of *σ*. The final selected parameter combination was *µ* = 2.0, *σ* = 0.80. This configuration provided the best qualitative agreement among the tested candidates, although discrepancies remained, particularly in the upper tail of the simulated count distribution.

Interestingly, converting the selected distributions to inferred mean underlying body numbers produced estimates of approximately 102 bodies for WT and 10 bodies for the mutant. These are nearly identical to the corresponding conventional stereological estimates of 101 and 10 bodies (Table II). The agreement in these summary estimates therefore contrasts sharply with the remaining disagreement between the simulated and experimental cross-section-count distributions. This result highlights an important distinction between body-size and body-number inference. While the selected body-size distributions reproduced the experimental observations comparatively well, the current spatial model was unable to reproduce the full body-count distributions despite yielding similar estimates of the underlying mean number of bodies.

## V. Discussion

The results of this study suggest that the value of Auto-AVS lies less in replacing conventional stereology than in testing the assumptions on which stereological inference depends. Across the validation and experimental fitting analyses, a consistent pattern emerged: inference of autophagic-body size was comparatively robust, whereas inference of body number was substantially more sensitive to assumptions concerning spatial organization and vacuole geometry.

For body-size estimation, Auto-AVS behaved similarly to both the analytical stereological calculation and the previously developed spherical R-based simulator. Although differences increased somewhat with greater variability in the underlying size distribution, the estimated equivalent cross-section radii remained comparatively stable across methods and body-count conditions. This behavior was also observed in the experimental fitting analysis. The final WT and mutant Auto-AVS size estimates were close to those obtained using conventional stereological calculations, and the selected simulated cross-section distributions reproduced the corresponding experimental distributions reasonably well. Taken together, these results indicate that the estimation of underlying body size is relatively insensitive to the additional spatial complexity introduced by the current Auto-AVS model.

Body-number inference, in contrast, showed considerably greater sensitivity to the underlying spatial model. In the validation experiments, Auto-AVS increasingly diverged from the analytical prediction for the fraction of generated bodies intersected by a virtual section as body count, mean body size, and body-size variability increased. This divergence was not mirrored to the same extent in the R-based simulation, suggesting that the discrepancy is associated with the explicit representation and packing of three-dimensional body populations rather than with the basic geometry of virtual sectioning alone.

The experimental fitting results reinforce this distinction. For both WT and mutant datasets, the final inferred mean body numbers were nearly identical to those obtained using conventional stereological calculations. However, the agreement in these summary estimates masked substantial differences between the simulated and experimental cross-section-count distributions. In particular, the WT simulations produced too many sections containing no visible bodies while also generating an overly heavy upper tail. This behavior is consistent with the compact clustering imposed by the current placement procedure: a virtual section may miss the cluster entirely, but when it intersects the cluster it can contain many bodies simultaneously. Thus, similar inferred mean body numbers do not necessarily imply that the simulated spatial organization is a faithful representation of the biological in vivo system.

This observation is important because conventional stereological approaches typically operate through summary relationships between observed cross-sections and inferred three-dimensional quantities. Auto-AVS provides an additional level of evaluation by allowing the full distribution of simulated observations to be compared with the experimental data. In the present study, this distribution-level comparison revealed deficiencies that would have been hidden if model assessment had relied only on mean estimates. The results therefore support the use of simulation-based stereology not only as an estimation tool, but also as a mechanism for testing whether the structural assumptions underlying an estimate are compatible with the observed data.

The CompuCell3D deformation experiments provide a second example of this role. Explicit deformation produced visibly non-spherical APB geometries, but the resulting changes in observed cross-section size and number were small, inconsistent, and not reproducible across the tested configurations. One initially significant difference in observed size was not reproduced in subsequent independent runs and disappeared when the runs were pooled. Under the present simulation parameters, therefore, the degree of deformation generated by CompuCell3D was not sufficient to measurably alter the section-level quantities used for inference.

This negative result should not be interpreted as evidence that body shape is universally irrelevant to stereological inference. The current experiments establish only that the specific degree and type of deformation produced under the tested settings did not produce a detectable change in the measured observables. More extensive deformation, different mechanical constraints, or alternative spatial arrangements could yield different results.

Several limitations of the current model are exposed directly by these analyses. Auto-AVS presently generates a compact APB cluster centered within the vacuole. The vacuole itself is represented using an idealized spherical geometry. The WT body-number mismatch suggests that the degree, consistency, or location of clustering in the simulation differs from that observed experimentally. Likewise, the geometry of the vacuole may influence the probability that a section intersects a given body population. These effects are likely to be particularly important for body-number inference because the observable count depends jointly on body abundance, body size, spatial arrangement, and the geometry of the containing vacuole.

Future work should therefore focus on relaxing these assumptions rather than simply increasing the resolution of the current parameter search. Potential extensions include alternative clustering models, off-center or membrane-proximal placement, non-spherical or experimentally informed vacuole geometries, and differently parameterized deformation models. Because Auto-AVS separates geometry generation, deformation, sectioning, and statistical evaluation into configurable stages, such hypotheses can be tested within the same framework and compared directly against experimental distributions.

Overall, the results demonstrate that simulation-based stereology can reveal where conventional estimates are robust and where they depend strongly on assumptions that are difficult to evaluate analytically. For autophagic-body size, the tested methods converged closely and produced simulated observations that agreed well with experiment. For body number, agreement in inferred means concealed substantial distributional mismatch. Auto-AVS therefore provides its greatest value as a framework for interrogating the assumptions behind three-dimensional inference from two-dimensional microscopy, rather than as a simple replacement for existing stereological calculations.

## VI. Conclusion

Auto-AVS provides a computational framework for testing the assumptions underlying stereological inference from TEM cross-sections of yeast autophagic bodies. Across validation and experimental fitting, body-size estimates were comparatively robust, while body-number inference was substantially more sensitive to spatial organization and vacuole geometry. CompuCell3D-based deformation did not produce a reproducible measurable effect under the tested settings. Overall, Auto-AVS is most useful not as a replacement for conventional stereology, but as a tool for identifying when apparently reasonable estimates are inconsistent with the full experimental distributions.

## VII. ACKNOWLEDGMENT

The authors thank Victor Tyler Jr. and Shaun Lee for their assistance with early-stage development and testing that helped inform the evolution of this work.

---

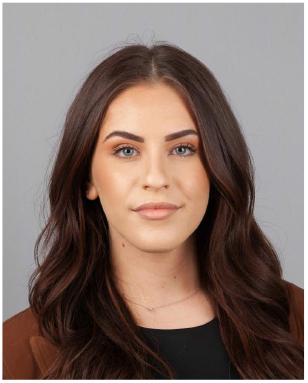

**Stela Bleta** is a graduate student in Computer Science at Eastern Michigan University, where she is currently pursuing her master’s degree. She earned her Bachelor’s degree in Computer Science from Eastern Michigan University in 2025. Her academic and research interests include artificial intelligence, machine learning, software development, and the use of computing technologies in education and healthcare.

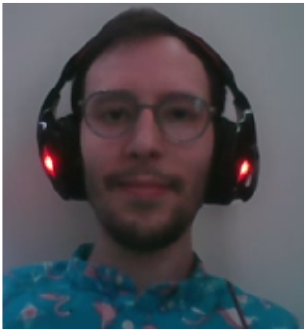

**Payton Dunning** graduated from Eastern Michigan University with a Bachelor’s in Biochemistry and Computer Science. At EMU he studied under Dr. Steven Backues investigating mechanisms and proteins related to the process of autophagy in baker’s yeast. Payton later attended the University of Michigan where he investigated the properties and structures of lysine methyltransferase enzymes in Bacillus bacteria, graduating with a Master’s degree in Biological Chemistry.

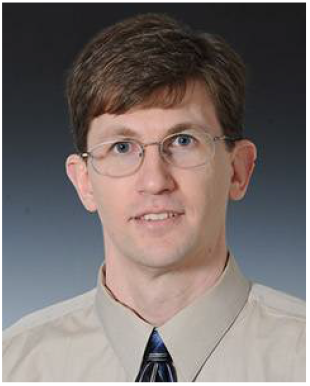

**Andrew Ross** is a Professor in the Department of Mathematics and Statistics at Eastern Michigan University. He earned his Ph.D. in Operations Research from the University of California, Berkeley. His research interests span applied mathematics, statistics, operations research, mathematical modeling, and interdisciplinary computational applications. His recent collaborative work includes the development and analysis of computational methods for studying autophagy and quantifying autophagic structures in transmission electron microscopy images.

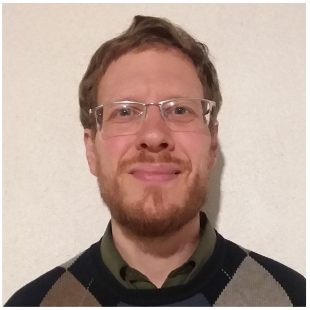

**Steven K. Backues** is a Biochemistry Professor in the School of Natural Sciences at Eastern Michigan University. He earned his PhD from the University of Wisconsin-Madison, studying membrane trafficking in plant cell growth, and did postdoctoral research on autophagy with Dr. Dan Klionsky at the University of Michigan. His current research uses baker’s yeast to understand the basic mechanisms of autophagy, a “self-eating” membrane-trafficking pathway that helps cells stay healthy and survive starvation.

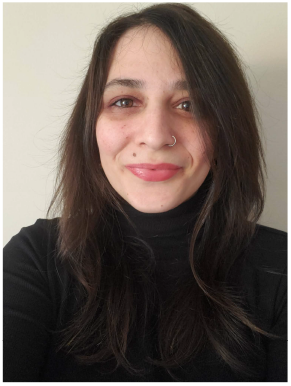

**Ourania Spantidi** is an Assistant Professor in the Computer Science Department at Eastern Michigan University and the director of the Embedded AI Systems Lab. She earned her Ph.D. from the Embedded Systems Software Lab in the School of Electrical, Computer, and Biomedical Engineering at Southern Illinois University. Her research interests lie in the area of applied artificial intelligence, human-computer interaction, approximate computing, and energy-efficient machine learning systems.

## Footnotes

1 https://github.com/eas-lab-emich/Auto-AVS

